# PAR2 signaling shapes microbial and metabolic remodeling along the gut–lung axis

**DOI:** 10.64898/2026.09.11.751027

**Authors:** Ritu Mann-Nüttel, Riya Jindal, Marie Armbruster, Shivani Mandal, Gang Zhou, Chelsey Blackman, Maryalle Tan, Harissios Vliagoftis, Paul Forsythe

## Abstract

**Introduction:** Protease activated receptor (PAR2) is a prominent sensor of environmental and microbial proteases and may serve as an important interface between the host epithelium and mucosal microbes. However, whether PAR2 helps shape microbial community structure and function at barrier surfaces remains unclear. To address this, we examined how PAR2 deficiency or activation affects microbial composition and metabolic potential across the gut lumen, airway lumen, and lung tissue in the context of exposure to protease-rich house dust mite allergen (HDM) exposure.

**Methods:** Wild-type and PAR2-deficient littermates of both sexes received a single intranasal challenge with phosphate-buffered saline, HDM extract, or a selective PAR2 agonist. Microbial communities from feces, bronchoalveolar lavage fluid (BALF), and lung tissue were profiled using 16S rRNA gene sequencing and PICRUSt2-based functional inference to assess compartment-specific taxonomic and metabolic responses.

**Results:** Alpha and beta diversity remained stable across all experimental groups, but distinct conditions resulted in compartment-specific remodeling of the microbial population. At baseline, PAR2 deficiency altered multiple genera in the gut and lung and shifted predicted pathways linked to amino-acid, lipid, and sulfur metabolism. HDM induced broad taxonomic and functional changes in the gut and lung tissue—including shifts in coenzyme A biosynthesis, reductive TCA activity, lysine fermentation, and nucleotide biosynthesis—while producing only limited taxonomic changes in BALF. PAR2 signaling accounted for a substantial portion of HDM-driven remodeling, and direct PAR2 activation reproduced many compartment-specific effects, including mucin-derived sugar degradation in the gut and suppression of nucleotide biosynthesis in the lung. Sex moderately modified microbial and metabolic responses, with males and females exhibiting divergent, condition-dependent functional biases across gut and lung.

**Conclusion:** These findings identify PAR2 as a mucosal niche-modifying receptor whose activation or loss reshapes microbial composition and metabolic potential along the gut–lung axis.

## 1. Introduction

Mucosal surfaces, such as the gut and lung, are continuously exposed to environmental proteases, allergens, and microbes. Microbial communities at these sites play a central role in shaping mucosal immunity and alterations in their composition, such as changes in the relative abundance of major phyla like *Bacteroidota* and *Bacillota*, have been linked to susceptibility to respiratory infections^1^ and asthma^2^. While drug exposures and disease conditions have been extensively examined for their impact on the microbiome, far less is known about how host gene expression shapes microbial composition and function. Pattern-recognition receptors such as Toll-Like Receptor (TLR) 4 exemplify how host sensing pathways can shape microbial communities while simultaneously being influenced by microbial products^3^. Similar principles likely apply to other mucosal receptors that detect environmental cues, yet their roles in regulating microbiome structure and function remain poorly defined. Protease-activated receptor-2 (PAR2), a major sensor of microbial^4^ and environmental proteases^5^, is one such receptor. PAR2 has well-established roles in allergic airway disease, with genetic and functional studies linking PAR2 to asthma susceptibility and severity^6,7^. PAR2-deficient mice exhibit reduced airway hyperresponsiveness and diminished inflammation following allergen challenge^7,8^. Conversely, PAR2 activation promotes allergic sensitization to co-inhaled antigens^9^ and amplifies allergen-driven airway inflammation^10^. Yet despite these well-defined immunological functions, the role of PAR2 in shaping mucosal microbiomes remains unknown, even though microbial communities rely heavily on protease-mediated mechanisms for nutrient acquisition^11^ and for communication with the host^12^.

In this study, we aimed to characterize the impact of PAR2 on the microbial community of the lung and gut. Microbial communities were profiled across feces (gut lumen), bronchoalveolar lavage fluid (BALF, airway lumen), and lung tissue to capture compartment-specific ecological responses in both female and male wild-type (WT) and PAR2 deficient mice. Microbiome changes were also assessed following a single acute intranasal challenge with either a selective PAR2 agonist or HDM extract; a physiologically relevant allergen rich in PAR2 activating proteases. By integrating 16S rRNA gene profiling with PICRUSt2-based these analyses were designed to define the contribution of PAR2 signaling to microbial community composition and metabolic potential across the gut–lung axis.

## Materials and Methods

### Experimental Setup

All animal procedures were approved by the University of Alberta Animal Care and Use Committee (AUP00000353) and conducted in accordance with institutional guidelines. Across four litters (L1–L4), both WT (F2rl1+/+) and PAR2^-/-^ (F2rl1tm1a/tm1a) C57/BL6 mice contributed to the study cohort (**Fig. S1**), allowing assessment of genotype- and sex-specific responses. Cage effects were minimized by combining litters at weaning. By 8 weeks of age, all mice— despite a two-day age offset in L2—had reached comparable developmental stages, and treatment groups contained balanced representation from each litter. Mice were housed in groups of five per ventilated cage, had ad libitum access to food (LabDiet 5L0D) and water (changed weekly), and were maintained on a 12-hour light–dark cycle at 22°C and 52.1% humidity. Mice were administered one of three intranasal stimuli—PBS, house dust mite (HDM) extract, or a PAR2-activating peptide (PAR2-AP)—and evaluated 18□h after exposure, a time point selected to capture early microbial and metabolic shifts along the gut–lung axis. Low-endotoxin HDM extract (XPB91D3A2.5; Stallergenes Greer, Lenoir, NC, USA) was reconstituted in endotoxin-free 1× PBS (HyClone, Cytiva, USA). A 25□µL intranasal dose containing 1□µg/µµL HDM (protein weight) was administered under 2–5% isoflurane anesthesia. For targeted PAR2 activation, mice received 25□µL of 100□µM PAR2-activating peptide (Synpeptide Co., Ltd., Shanghai, China) intranasally, delivered under the same anesthesia conditions. Microbial communities were characterized across three mucosal compartments. Fecal samples reflected distal intestinal microbiota, and BALF and lung tissue represented two different techniques to assess airway-associated communities.

**Figure S1. Experimental design and sample collection workflow.**

### Feces Collection and processing

Fecal pellets were obtained directly from the distal intestine and deep-freezed in liquid nitrogen and maintained in at −80□°C freezer until further processing. BALF was collected by instilling and retrieving PBS through the trachea. Lung tissue was sampled by excising the left lung lobe and mechanically disrupting the tissue after taking BALF. For genomic DNA isolation, stool or tissue was transfer into tubes with freshly prepared buffer (proteinase K, RNase, buffer RTL from the Qiagen Dneasy® Blood & Tissue Kit). Samples were homogenized by vortex for 1 minute and incubated at 56°C 75 min. on a shaker with intermittent vortex every 15 min. 16S bacterial genomic DNA was then isolated using the Dneasy® Blood & Tissue Kit (Cat: 69506, Qiagen) following the vendor instructions. Bronchoalveolar lavage (BAL) samples were incubated with Biofluid and Cell Buffer and Proteinase K at 55□°C for 90□min on a shaker set to maximum speed. Samples were briefly vortexed for 15□s every 20□min to ensure thorough lysis. Following enzymatic digestion, genomic DNA was isolated using the Quick-DNA Miniprep Plus Kit (Zymo Research, D4068) according to the manufacturer’s instructions.

### 16S DNA sequencing of microbiome

Quality and quantity were checked using the NanoDrop® ND-1000 Spectrophotometer (NanoDrop® Technologies). Samples were submitted to Novogene Corporation Inc. (Sacramento, CA, USA) for 16S rRNA gene amplicon sequencing targeting the V3–V4 region using the Illumina NovaSeq 6000 platform, generating approximately 30,000 paired⍰end reads per sample. Raw FASTQ files were processed in⍰house. Primer trimming was performed using Cutadapt and read quality was assessed using FastQC^13^. Reads were filtered at phred ≥ 30. Amplicon sequence variants (ASV) were inferred using DADA2^14^ with error⍰model learning, dereplication, paired⍰end merging, and chimera removal. Taxonomic assignment was performed using the Genome Taxonomy Database (GTDB) reference database, release 09⍰RS220. ASV tables were rarefied to a depth of 47,055 reads prior to diversity analyses. Beta⍰diversity metrics (Bray–Curtis), PERMANOVA tests, visualization of taxonomic composition and alpha diversity metrics were computed in QIIME2^15^ 2023.9. Functional pathway predictions were generated using PICRUSt2^16^. All sequencing data have been deposited in the NCBI BioProject database under accession number PRJNA1469202.

### Statistical analysis

Alpha Diversity differences were assessed using pairwise Wilcoxon rank-sum tests. P-values were adjusted for multiple comparisons using the Benjamini–Hochberg false discovery rate (FDR) method. Beta diversity was assessed using Bray–Curtis dissimilarity calculated from genus-level relative abundance matrices. Ordination was performed using principal coordinates analysis (PCoA) to visualize community structure. To test for differences in overall microbial composition, PERMANOVA (adonis2, 999 permutations) was applied with the following model for each compartment: Bray–Curtis ∼ Sex + Genotype + Treatment. This approach evaluates the main effect of sex while adjusting for genotype and treatment, avoiding over-stratification of the dataset. Homogeneity of group dispersions was assessed using PERMDISP to ensure that significant PERMANOVA results were not driven by differences in variance. To identify genera differing between experimental groups, absolute genus-level abundances were compared using Wilcoxon rank-sum tests for each pairwise contrast of interest. Because these analyses were intended to provide descriptive, exploratory insight into sex-, genotype-, and treatment-associated shifts, no multiple-testing correction was applied. All statistical tests were performed separately for gut, lung, and BALF compartments. The test used is specified for each Figure legend. All bar graphs show mean and error bars with SEM. Copilot (GenAI) was employed to generate the R code used to visualize data (**Fig. 1, 2, 3**) and Figures were further modified in Inkscape (v.1.4.3).

**Figure 1.**
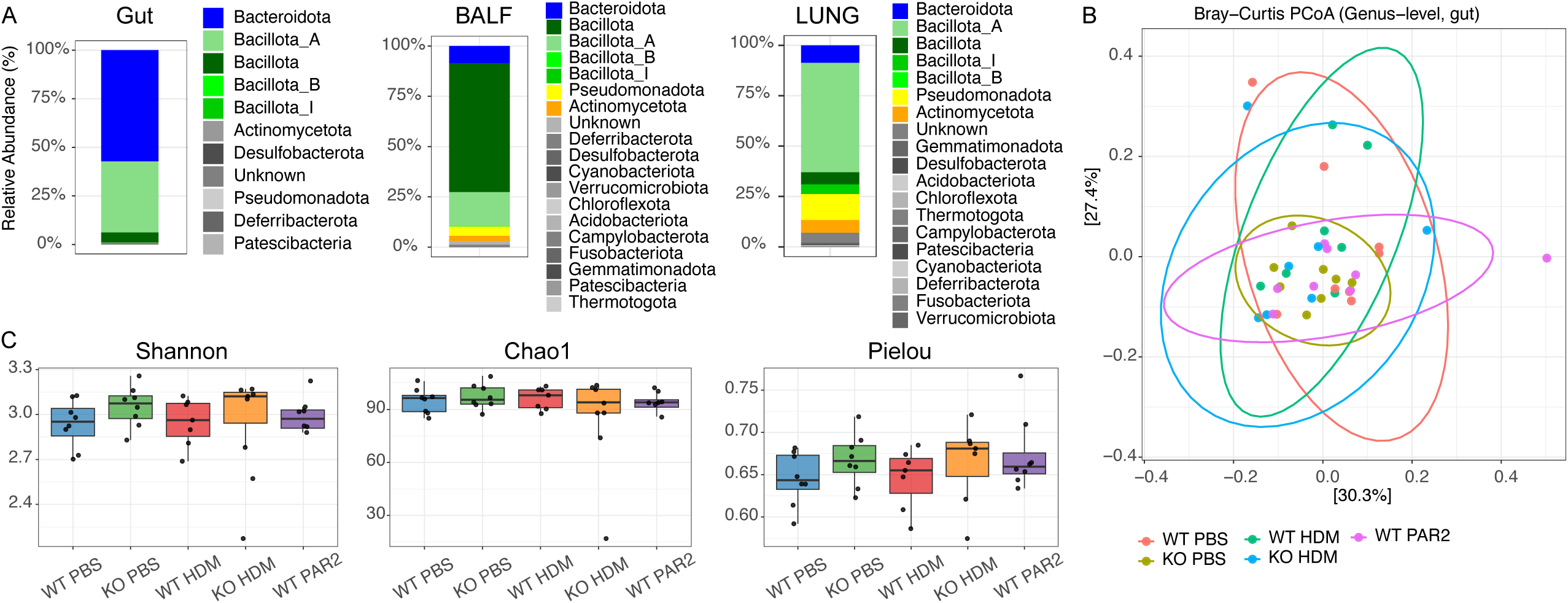
Phylum-level composition and diversity across gut, BALF, and lung microbiomes. **A** Stacked bar plots showing phylum-level community composition in gut, BALF, and lung samples from the wild-type PBS group. **B** Principal coordinates analysis (PCoA) of Bray–Curtis dissimilarities illustrating overall community structure across the three compartments. **C** Alpha-diversity metrics (Shannon, Chao1, Pielou’s evenness) displayed for each compartment and experimental group.

**Figure 2.**
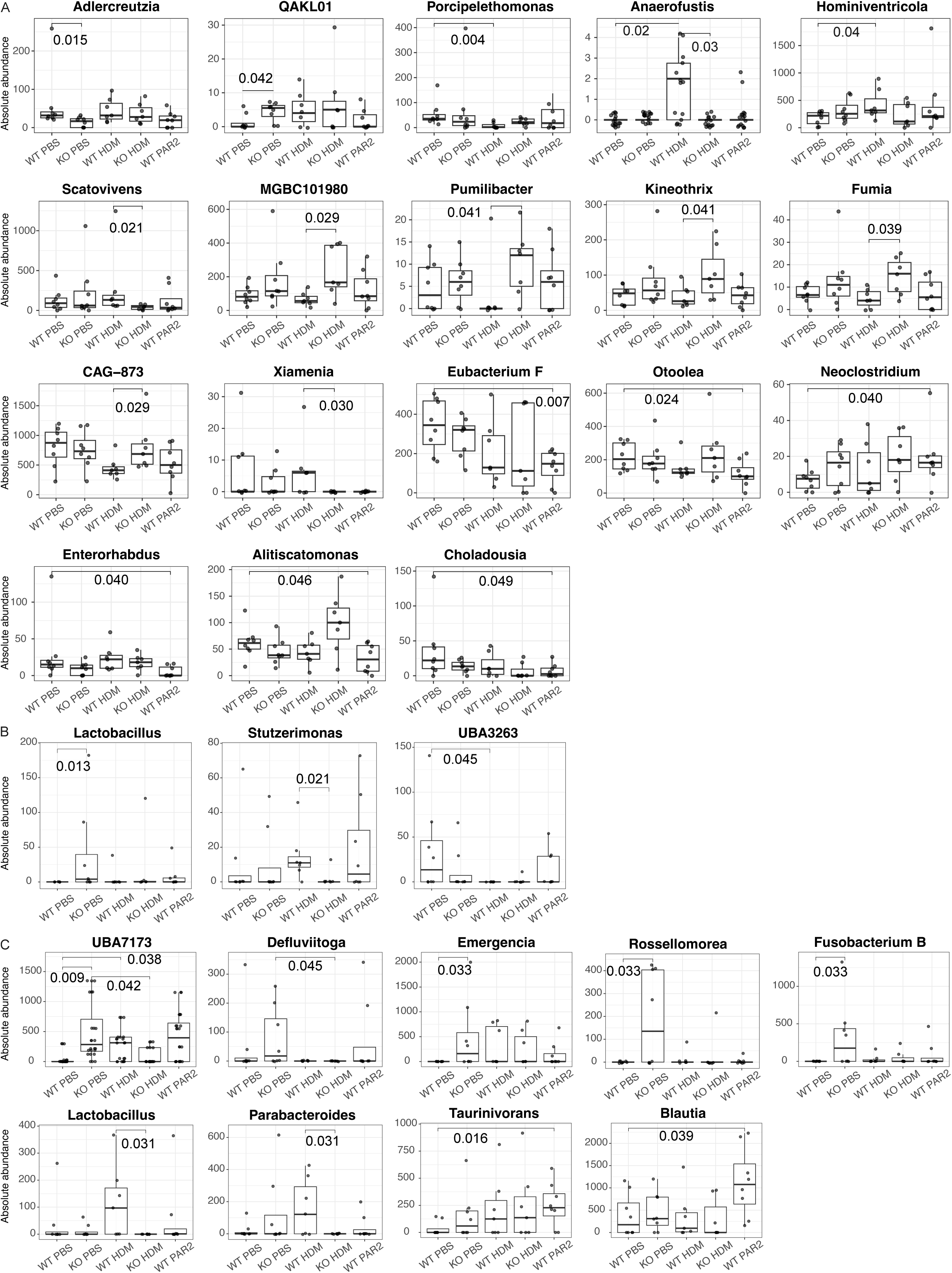
Absolute abundance profiles across gut, BALF, and lung. Gut absolute abundance values across WT, PAR2^-/-^, WT + HDM, PAR2^-/-^ + HDM, and WT + PAR2 agonist groups for gut (**A**), BALF (**B**) and lung (**C**). Group comparisons were assessed using Wilcoxon rank-sum tests.

**Figure 3.**
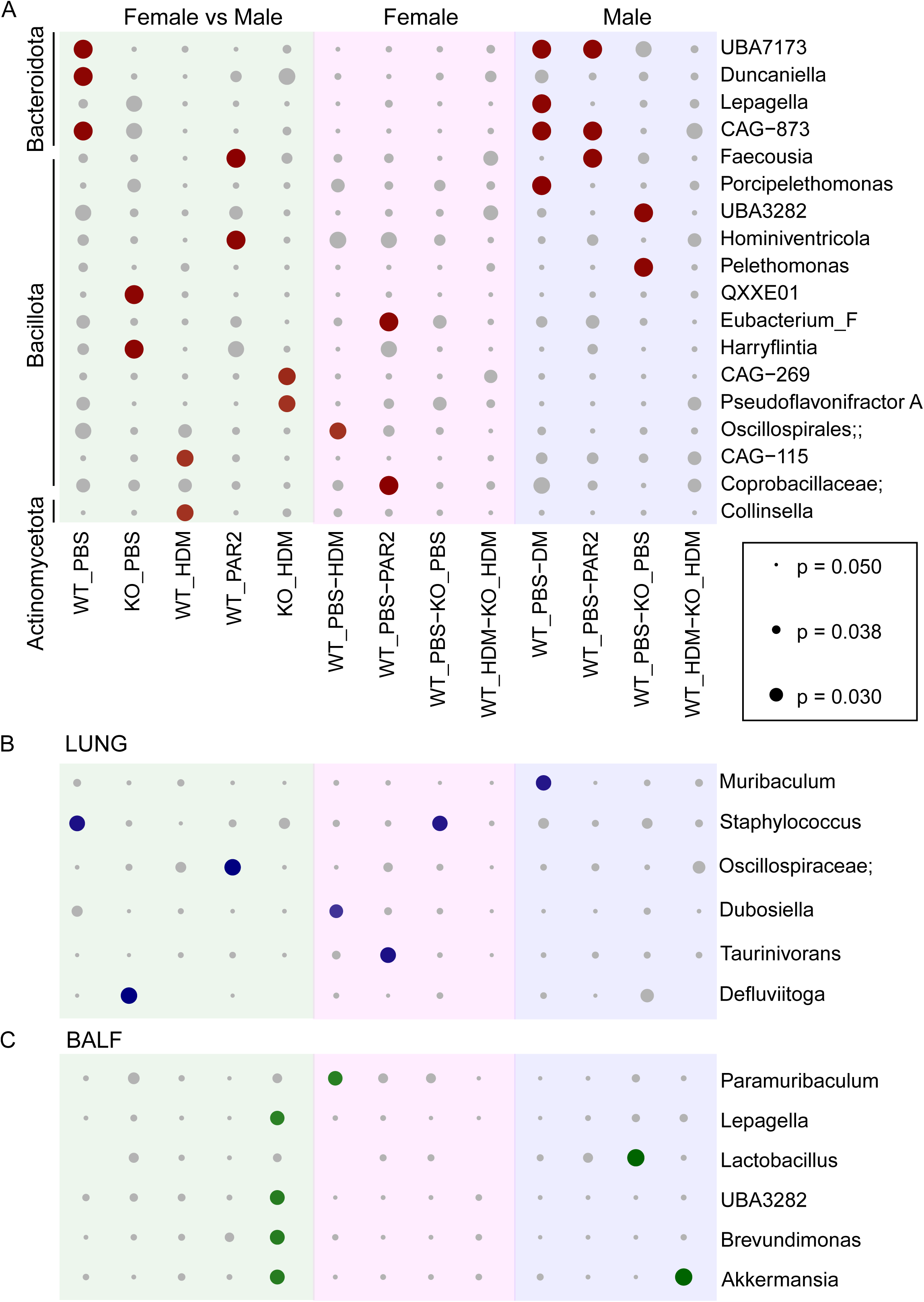
Genus-level comparisons of sex-associated and within-sex differences across gut, lung, and BALF. Dot plots display genus-level absolute abundance comparisons for three predefined contrasts: female vs male (green), within-female (pink), and within-male (blue) in gut (**A**), lung (**B**) and BALF (**C**). Significant comparisons are shown as colored dots, while non-significant comparisons (p > 0.05) appear in grey. Dot size reflects the corresponding p-value, with smaller p-values represented by larger dots. All comparisons were assessed using Wilcoxon rank-sum tests.

## 2. Results

### 2.1 Phylum-level community structure and diversity across gut, BALF, and lung samples

Across all three compartments—gut, BALF, and lung tissue—the microbial communities were dominated by a small number of bacterial phyla (**Fig. 1A** showing WT samples), with compartment-specific differences in overall composition but no significant diversity shifts across experimental groups (**Fig. 1B, C**). In the gut, the major phyla were *Bacteroidota* (57.2% ± 4.0%) and *Bacillota A* (36.4% ± 3.8%), followed by *Bacillota* (5.3% ± 3.8%), with minor contributions from *Actinomycetota* (0.6% ± 0.3% SE) and *Pseudomonadota* (0.08% ± 0.03%). In BALF, the community was dominated by *Bacillota* (mean 64.0% ± 13.8% SE) and *Bacillota A* (17.1% ± 8.3%), with additional contributions from *Bacteroidota* (8.5% ± 4.6%), *Pseudomonadota* (4.6% ± 2.6%), and *Actinomycetota* (2.8% ± 1.1%). In lung tissue, the community was enriched for *Bacillota A* (54.4% ± 4.3%), *Pseudomonadota* (12.8% ± 6.4%), *Bacteroidota* (8.6% ± 1.5%), and *Actinomycetota* (6.4% ± 1.4%), with additional contributions from *Bacillota* (6.0% ± 2.7%) and *Bacillota I* (4.7% ± 0.8%). Details of low abundance phyla can be found in **Table S3.**

### 2.2 Beta and alpha diversity remained stable across genotype and treatment within each compartment

Principal coordinates analysis (PCoA) based on Bray–Curtis dissimilarities for beta diversity showed clear separation between gut, BALF, and lung communities, reflecting strong compartment-specific microbial signatures. However, within each compartment, neither genotype nor treatment produced significant shifts in overall community structure (**Fig. 1B** showing gut). This lack of clustering is notable given that HDM activates PAR2^17^ alongside multiple other pattern-recognition pathways, whereas the PAR2 agonist selectively engages PAR2 without triggering broader receptor networks. Despite these mechanistic differences, pairwise PERMANOVA tests confirmed the absence of group-level differences in community composition across all comparisons (**Table S4**). In the gut, all pairwise contrasts showed small effect sizes (R² = 0.03–0.10) and non-significant p-values (p > 0.20). Lung communities exhibited similarly low effect sizes (R² = 0.04–0.07) with no significant differences (p > 0.47), and BALF showed the smallest effects overall (R² = 0.007–0.023) with uniformly high p-values (p = 0.77–0.98). Similarly, across all three compartments alpha diversity metrics were comparable between genotype and treatment groups (**Fig. 1C** showing gut; **Table S4**). In the gut, pairwise Wilcoxon tests showed no significant differences in Shannon diversity, Chao1 richness, or Pielou evenness across any comparison (Gut, p > 0.19; Lung, p > 0.15; BALF, p > 0.38). Together, these results demonstrate that within-sample diversity remained stable across compartments and was not influenced by PAR2 status, HDM exposure, or PAR2 agonism, consistent with the absence of broad community-level restructuring.

### 2.3 PAR2 deficiency is associated with taxonomic and functional changes to the lung and gut microbial communities

Given the absence of broad diversity shifts, we next evaluated genus-level taxonomic differences. and assessed predicted functional profiles using PICRUSt2 (**Table S5**) to determine whether treatment or genotype influenced microbial metabolic potential despite stable alpha and beta diversity.

At baseline (PBS treatment), PAR2 deficiency was associated with significant shifts in both gut and airway microbial communities (**Fig. 2**). In fecal samples, *Adlercreutzia* and *QAKL01* were reduced in PAR2^-/-^ relative to WT (p = 0.015 and p = 0.042, respectively), indicating that intact PAR2 signaling contributes to maintenance of these specific gut taxa (Fig. 3A). PICRUSt2 analysis identified one significantly altered functional pathway in the gut (diaminopimelate/lysine biosynthesis), which showed higher predicted relative abundance in PAR2^-/-^ compared with WT (FDR = 0.0419; **Table S5**). Higher or lower predicted abundance reflects the relative contribution of the community to that pathway’s gene content, meaning a higher value indicates a greater predicted functional potential for that pathway. In BALF, *Lactobacillus* differed between WT and PAR2^-/-^ (p = 0.013), suggesting a parallel, tissue-specific baseline effect of PAR2 on the airway-associated microbiota (Fig. 3B). No significant PICRUSt2 pathways were detected in BALF at baseline. In lung tissue, several genera also showed genotype-linked differences under PBS conditions: *UBA7173* (*Muribaculaceae*) differed between WT and PAR2^-/-^ (p = 0.009), and *Emergencia*, *Rossellomorea*, and *Fusobacterium B* were likewise altered in KO PBS compared with WT (all p = 0.033, Fig. 3C). These taxonomic differences were accompanied by multiple significant PICRUSt2-inferred functional shifts (**Table S5**). PAR2^-/-^lungs showed higher predicted abundance of palmitate biosynthesis II (FDR = 0.0130) and the superpathway of sulfolactate degradation (FDR = 0.0480), whereas WT lungs showed higher predicted abundance of several chlorophyllide biosynthesis pathways (CHLOROPHYLL-SYN; PWY-5531; PWY-7159; all FDR ≈ 0.034–0.036). Together, these results indicate that PAR2 influences mucosal microbial composition and predicted functional potential at baseline across compartments, with both taxonomic and pathway-level differences that are largely tissue specific.

#### 2.3.1 HDM specific microbial responses in WT mice

A single acute intranasal HDM exposure produced clear but compartment-specific microbial shifts in WT mice, with multiple genus-level and pathway-level changes in the gut and lung tissue but a more limited response detected in BALF (**Fig. 2, Table S5**). In fecal samples, HDM treatment in WT mice altered the abundances of *Porcipelethomonas*, *Anaerofustis*, and *Hominiventricola* (all p = 0.03), consistent with HDM-driven remodeling of multiple gut lineages. These genus-level shifts were accompanied by two significant PICRUSt2-inferred pathway differences: WT + HDM samples showed lower predicted abundance of the coenzyme A biosynthesis pathway (FDR = 0.0174) and higher predicted abundance of the reductive citric acid cycle (FDR = 0.043) relative to WT. In BALF, HDM exposure in WT animals was associated with a significant change in *UBA3263* (p = 0.045), indicating that microbiome sampled by BALF is also influenced by allergen challenge. No significant PICRUSt2 pathway changes were detected in BALF microbiome. In lung tissue, HDM exposure produced significant differences for several genera, including a WT versus WT + HDM difference for *UBA7173* (p = 0.038). These taxonomic changes coincided with multiple significant PICRUSt2-inferred pathway shifts in the lung microbiome (**Table S5**). WT + HDM lungs showed higher predicted abundance of L-lysine fermentation to acetate and butanoate pathway (FDR = 0.0188), whereas WT lung microbiome showed higher predicted abundance of several de novo purine and pyrimidine biosynthesis superpathways (DENOVOPURINE2-PWY; PWY-841; PWY-6125; PWY0-162; PWY-7184; PWY-7196; PWY-7228; all FDR ≈ 0.024–0.045). Together, these findings show that HDM elicits a broad microbial response in the gut and lung tissue—affecting both taxa and predicted functions—while the BALF samples exhibit a more restricted, primarily taxonomic shift, highlighting distinct sensitivities of mucosal compartments to allergen exposure.

#### 2.3.2 PAR2 dependent microbial responses to HDM

Comparing WT + HDM with PAR2^-/-^ + HDM revealed that PAR2 signaling shapes a substantial portion of the HDM-induced gut response, whereas in the respiratory tract PAR2 dependence was more compartment-specific, with few genus-level changes detected in BALF microbiome and a broader set of differences in the lung tissue samples (**Fig. 2, Table S5**). In the gut, multiple genera that changed with HDM in WT mice failed to do so in PAR2-deficient animals; notable PAR2-dependent taxa included *Scatovivens* (p = 0.021), *MGBC101980*, *Pumilibacter*, *Kineothrix*, *Fumia*, *CAG-873*, and *Xiamenia* (all p = 0.030). No significant PICRUSt2-inferred pathways were detected in the gut for this comparison. In BALF, only *Stutzerimonas* showed a PAR2-dependent HDM response (WT + HDM vs PAR2^-/-^+ HDM, p = 0.029), and no significant PICRUSt2 pathways were identified. In lung tissue, several OTUs exhibited PAR2-linked differences. For example, *UBA7173* and *Defluviitoga* differed between PAR2^-/-^ and PAR2^-/-^ + HDM (p = 0.042 and p = 0.045, respectively), and importantly, *Lactobacillus* and *Parabacteroides* were altered between WT + HDM and PAR2^-/-^ + HDM (both p = 0.031), indicating that PAR2 status modulates how both BALF and lung tissue communities respond to allergen exposure. These genus-level differences were accompanied by multiple significant PICRUSt2-inferred pathway shifts in the lung (**Table S5**). WT + HDM lungs showed higher predicted abundance of the superpathway of geranylgeranyl-diphosphate biosynthesis I (FDR = 0.012), the mevalonate pathway I (FDR = 0.012), the peptidoglycan biosynthesis V (FDR = 0.021), L-lysine biosynthesis II (FDR = 0.027), peptidoglycan biosynthesis II (FDR = 0.028), L-phenylalanine biosynthesis (FDR = 0.0310), L-tyrosine biosynthesis (FDR = 0.031), L-histidine degradation I (FDR = 0.037), and L-isoleucine and L-valine biosynthesis (both FDR = 0.048). Overall, PAR2 dependence of HDM responses was strongest and most widespread in the gut at the genus level, minimal in the airway lumen, and selective but functionally extensive in the lung tissue, where numerous pathway-level differences reflected substantial shifts in predicted metabolic potential.

#### 2.3.3 PAR2 agonist induced microbial shifts

Direct PAR2 activation produced a compartmental pattern that resembled the HDM response—broad changes in the gut, minimal alterations in BALF, and functional remodeling in lung tissue—but with markedly different magnitude and biological drivers (**Fig.2**, **Table S5**). In fecal samples, PAR2 agonism in WT mice altered multiple genera—including *Eubacterium F*, *Otoolea*, *Neoclostridium*, *Enterorhabdus*, *Alitiscatomonas*, and *Choladousia* (all p = 0.030)—demonstrating that PAR2 activation alone is sufficient to remodel gut community structure. These genus-level changes were accompanied by several significant PICRUSt2-inferred pathway differences in the gut. WT + PAR2 agonist samples showed higher predicted abundance of the superpathway of L-rhamnose and L-fucose degradation (FDR = 0.007), the L-fucose degradation pathway (FDR = 0.023), and the superpathway of L-rhamnose degradation (FDR = 0.031), and lower predicted abundance of the superpathway of adenosylcobalamin biosynthesis from cobinamide-GMP (FDR = 0.020) and coenzyme A biosynthesis (FDR = 0.046). In BALF, no genera reached significance in the WT + PAR2 versus WT comparison, and no significant PICRUSt2 pathways were detected, suggesting that the airway lumen is less responsive to direct PAR2 stimulation. In lung tissue, PAR2 agonism (WT + PAR2) was associated with changes in *Taurinivorans* and *Blautia* when compared with WT (p = 0.016 and p = 0.039, respectively), indicating that PAR2 activation can influence specific lung tissue taxa even when BALF shows no global response. These taxonomic differences coincided with multiple significant PICRUSt2-inferred pathway shifts in the lung. WT + PAR2 agonist lungs showed lower predicted abundance of the superpathway of purine nucleotides de novo biosynthesis II (FDR = 0.022), pyridoxal-5′-phosphate biosynthesis I (FDR = 0.025), the superpathway of guanosine nucleotides de novo biosynthesis II (FDR = 0.026), the superpathway of purine nucleotides de novo biosynthesis I (FDR = 0.027), the superpathway of pyrimidine ribonucleotides de novo biosynthesis (FDR = 0.036), pyrimidine deoxyribonucleotide biosynthesis I (FDR = 0.042), the superpathway of pyrimidine ribonucleosides salvage (FDR = 0.043), the superpathway of guanosine nucleotides de novo biosynthesis I (FDR = 0.048), the superpathway of histidine, purine, and pyrimidine biosynthesis (FDR = 0.045), and the superpathway of pyridoxal-5′-phosphate biosynthesis and salvage (FDR = 0.049). In contrast, WT + PAR2 agonist lungs showed higher predicted abundance of myo-, chiro-, and scillo-inositol degradation (FDR = 0.029), TCA cycle VII (acetate-producers) (FDR = 0.044), and the methylaspartate cycle (FDR = 0.046).

### 2.4 Males and female differences in gut and lung microbial and metabolic profiles

#### 2.4.1 Sex differences in Alpha Diversity

Alpha-diversity was stable across all compartments. Within each sex, no treatment or genotype produced significant differences in Shannon, Chao1, or Pielou values in the gut, lung, or BALF (all p > 0.11; **Table S6**). Female–male comparisons likewise showed no consistent differences in any compartment (all p > 0.21), aside from a single uncorrected Shannon contrast in the PAR2 agonist group in the lung (p = 0.03). Overall, richness and evenness remained comparable across sexes, treatments, and genotypes.

#### 2.4.2 Female vs male differences across compartments

Sex-associated differences occurred in all three compartments (**Fig.□3, Table S7**), strongest in the gut, followed by lung tissue, with fewer but distinct differences in BALF. Across tissues, the most consistently sex-responsive groups were *Muribaculaceae* (*Bacteroidota*) and Clostridia-associated *Bacillota A*, with additional contributions from *Actinomycetota*, *Thermotogota*, *Desulfobacterota*, and *Verrucomicrobiota*. In the gut, baseline female–male differences involved several *Muribaculaceae* genera, including *UBA7173* (p□=□0.03), *Duncaniella* (p = 0.03), and *CAG-873* (p□=□0.03). Under PAR2 stimulation, WT females and males differed in *Faecousia* (p□=□0.029) and *Hominiventricola* (p□=□0.030). In PAR2^-/-^ mice, baseline differences were detected for *QXXE01* (p□=□0.03) and *Harryflintia* (p□=□0.03). HDM exposure introduced additional sex-associated shifts in *CAG-269* (p□=□0.04), *Pseudoflavonifractor A* (p□=□0.04), CAG-115 (p□=□0.04), and *Collinsella* (p□=□0.04). In the lung, sex differences also spanned multiple phyla. WT + PAR2-treated mice differed in an unclassified Oscillospiraceae genus (p□=□0.02). Baseline PAR2^-/-^ comparisons showed differences in *Defluviitoga* (p□=□0.02). Additional contrasts involved *Staphylococcus* (p□=□0.0265), *Muribaculum* (p□=□0.03), and *Dubosiella* (p□=□0.049). In BALF, sex differences were fewer but phylogenetically diverse. Baseline contrasts identified *Lactobacillus* (p□=□0.02). Under HDM exposure in PAR2^-/-^ mice, sex differences occurred in *UBA3282* (p□=□0.04), *Brevundimonas* (p□=□0.0436), *Akkermansia* (p□=□0.04), and *Lepagella* (p□=□0.049).

### 2.5 Sex-specific PICRUST2-inferred functional differences

To determine whether functional microbial responses to PAR2 signaling and allergen exposure differ between females and males, we next performed sex-stratified PICRUSt2 analyses (**Table S8**). Across all conditions, sex-dependent functional differences were detected in the gut and lung, whereas no significant pathway-level differences were observed in BALF. For clarity, pathways described as “male-biased” or “female-biased” refer to those with greater predicted functional abundance in males or females, respectively. In WT mice, the gut showed 20 significant pathways. Most were male-biased (FDR□=□0.00034–0.036), involving cofactor, nucleotide, and amino acid biosynthesis, while a smaller subset was female-biased (FDR□=□0.007–0.049), primarily linked to carbohydrate degradation. In the lung, 9 pathways differed between sexes. Most were female-biased (FDR□=□0.007–0.047), spanning central metabolic and cofactor-related functions, with one male-biased degradation pathway (FDR□=□0.046). In PAR2^-/-^mice, the gut showed 10 significant pathways, most of which were male-biased (FDR□=□0.012–0.046), including cobalamin-associated and tetrapyrrole biosynthetic functions. Female-biased pathways were limited to fatty acid elongation and glycolysis (FDR□=□0.024–0.044). The lung also showed 10 pathways differing between sexes. Several central carbon and amino acid pathways were female-biased (FDR□=□0.014–0.040), whereas multiple biosynthetic and degradation pathways were male-biased (FDR□=□0.023– 0.049). In WT + HDM mice, the gut showed 3 significant pathways, all male-biased (FDR□=□0.019–0.048), primarily involving carbohydrate-degradation functions. The lung displayed 22 pathways, the largest sex-associated shift observed. Quinone-related and energy-linked pathways were strongly female-biased (FDR□=□0.0026–0.0157), whereas fermentation and amino acid biosynthesis pathways were male-biased (FDR□=□0.0067– 0.023). In WT + PAR2 mice, the gut showed 1 significant pathway, which was male-biased (FDR□=□0.024). The lung showed 4 pathways, including a female-biased nucleotide biosynthesis pathway (FDR□=□0.0186) and male-biased siderophore, aromatic-compound degradation, and carbohydrate-shunt pathways (FDR□=□0.0109–0.049). Together, these findings indicate that while sex influences similar metabolic themes across mucosal sites, the direction and context of these effects are highly compartment- and condition-dependent.

## 3. Discussion

This study provides the first analysis of how PAR2 signaling shapes microbial communities and their predicted metabolic activity across gut and lung under homeostatic conditions, acute allergen exposure, and direct PAR2 activation. Although alpha and beta diversity remained stable across all experimental groups, we identified robust compartment specific taxonomic and functional differences that were PAR2 dependent, and in some cases sex dependent. Baseline PAR2 deficiency altered both microbial composition and microbiome-derived predicted metabolic potential in the gut, BALF, and lung tissue. HDM exposure elicited broad microbial shifts in the gut and lung tissue, yet these microbiome-associated changes were poorly reflected in BALF. PAR2 signaling accounted for a substantial portion of these HDM-induced effects. Direct PAR2 agonism produced widespread microbial remodeling in the gut and selective microbial functional shifts in the lung, again with minimal effects detected in BALF. Finally, both taxonomic composition and microbiome-inferred predicted metabolic capacity differed between females and males, with the gut showing the most pronounced sex effects.

The phylum-level profiles in gut and lung tissue closely resemble those reported in a published dataset from C57BL/6 mice^18^, with *Bacillota*, *Bacteroidota*, *Actinomycetota*, and *Pseudomonadota* all falling within roughly five percent of the proportions typically illustrated for this strain. This similarity indicates that both compartments harbor relatively stable, tissue-associated microbial communities that are conserved across studies. In contrast, the BALF microbiome showed a markedly different phylum-level distribution, characterized by higher *Bacillota* and lower *Pseudomonadota* and *Actinomycetota* than commonly depicted in the literature^18^. This divergence reflects the fact BALF represents material recoverable from the airspaces and airway lumen, whereas tissue includes microbes associated with airway surfaces, parenchyma, and any adherent material retained in the lung. BALF microbial composition can be affected by lavage efficiency dilution and recovery efficiency and has a higher contamination risk than lung tissue. These methodological and ecological factors likely explain why gut and lung tissue align closely with published murine patterns, while BALF displays greater variability. These findings also support previous studies identifying lung tissue as a more robust choice for lung microbiome profiling^19^.

The absence of alpha and beta diversity changes across genotype and treatment groups indicates that PAR2 signaling and acute allergen exposure do not globally restructure microbial communities. Instead, the effects of PAR2 and HDM were subtle, targeted, and compartment specific, emerging at the level of individual genera and microbiome-inferred metabolic pathways. This pattern aligns with the concept that mucosal microbiomes are resilient at the community level but responsive at the functional and lineage-specific level, particularly under acute perturbation^20,21^ Given the role of PAR2 in sensing microbial and environment proteases, the targeted nature of these shifts is consistent with PAR2 acting as a local niche-modifying receptor rather than a global determinant of microbial diversity. Cuesta et□al. previously demonstrated that loss of the pattern-recognition receptor TLR4 alters gut microbial structure^3^, with TLR4^-/-^ mice showing reduced *Bacteroidota*, increased *Firmicutes* (*Bacillota*), and decreased *Actinobacteriota* (*Actinomycetota*). Remarkably, our PAR2^-/-^ mice exhibit the same directional shifts across these major phyla (**Table S2**), despite PAR2 sensing a very different class of environmental cues. One difference in the taxa shared between the two analyses was the behavior of *Desulfobacterota*, which increased in PAR2^-/-^ mice but decreased in the TLR4^-/-^ animals. Taken together, these parallels suggest that mucosal receptors—even those outside classical pattern-recognition families—may share common principles by which host sensing pathways shape microbial community structure. Across sexes combined, we observed 18 genus-level changes in the gut (14 occurring in *Bacillota* phylum, 3 in *Actinomycetota*, 1 in *Bacteroidota*), 3 in BALF (1 *Bacillota*, 1 *Bacteroidota*, 1 *Pseudomonadota*), and 9 in lung tissue (4 in *Bacillota*, 2 in *Bacteroidota*, 1 in *Thermotogota*, *Fusobacteriota* and *Desulfobacterota* each), underscoring the compartment-specific nature of microbial responses to PAR2 deficiency and HDM exposure. Although HDM was delivered intranasally and the challenge lasted only 18 hours, the larger compositional shifts in the gut than in airway samples are plausible: rapid innate and systemic responses to airway antigen (cytokines, acute-phase mediators, altered mucosal antibody trafficking) and possible swallowing of allergen-laden secretions can reach the intestine within hours and alter local niches. The protease load of HDM may also directly modify gut microbial nutrient landscapes by degrading host mucins and releasing glycan substrates that favor specific taxa. The far greater microbial biomass of the gut makes short-latency changes easier to detect than in low-biomass airway samples. Taxonomic shifts further occurred across a wide range of absolute abundances. In the gut, 8 of the 18 significantly altered OTUs had fewer than 50 absolute counts, whereas 10 OTUs showed substantially higher abundances, indicating that both dominant and low-abundance taxa contributed to the observed differences. BALF exhibited greater inter-mouse variability, but OTU abundances were generally below 100 counts, consistent with the low-biomass nature of airway lavage samples. In lung tissue, all 9 significantly altered OTUs exceeded 50 counts in at least one condition, reflecting a more stable and higher-biomass mucosal environment. We included both high- and low-abundance OTUs in our analysis because low-abundance taxa can exert disproportionate functional effects=. Statistically, we report non–multiple-testing-corrected p-values for genus-level comparisons. Although each compartment contained many detectable genera (129 in gut; 278 in lung and BALF; **Table S1**), strict multiple-testing correction would have removed biologically meaningful signals, particularly in low-biomass or high-variance contexts such as BALF. Our aim was exploratory identification of taxa contributing to PAR2-dependent shifts, with interpretation supported by microbial functional evidence from PICRUSt2. The gut exhibited 8 significant PICRUSt2-inferred pathways, whereas lung tissue showed 36 significant pathways, despite having fewer genus-level changes. BALF showed no significant pathways, consistent with its low biomass and limited number of responsive taxa. These findings demonstrate that the number of genera that change does not predict the magnitude of inferred microbiome-functional impact, as even a single OTU can influence multiple pathways, and coordinated shifts among several low-abundance taxa can reshape predicted metabolic outputs.

Across compartments, PICRUSt2-inferred pathways revealed that PAR2 signaling influences distinct aspects of microbial metabolism with potential consequences for mucosal physiology. In the gut, elevated diaminopimelate/lysine biosynthesis in PAR2^-/-^ mice suggests altered bacterial cell-wall metabolism and amino-acid turnover, processes that can affect microbial growth dynamics and the production of metabolites that shape epithelial integrity and immunological homeostasis. HDM exposure further shifted gut microbial function toward enhanced reductive TCA cycle activity and reduced coenzyme A biosynthesis, indicating changes in anaerobic energy metabolism and cofactor availability. PAR2 agonism produced yet another pattern, with increased rhamnose and fucose degradation—pathways linked to mucin-derived carbohydrate utilization—alongside reduced cobalamin and coenzyme A biosynthesis, suggesting altered microbial nutrient processing during direct receptor activation.

In lung tissue microbiome, functional shifts were more extensive and pointed toward changes in microbial lipid, amino-acid, and nucleotide metabolism. Increased palmitate biosynthesis and sulfolactate degradation in PAR2^-/-^ lungs indicate altered microbial lipid production and sulfur compound turnover, both of which can influence local redox balance and epithelial–microbial interactions^22^. HDM exposure in WT mice enhanced lysine fermentation to short-chain fatty acids, while reducing multiple de novo purine and pyrimidine biosynthesis pathways, suggesting a shift from proliferative to fermentative metabolic states within the lung-associated microbiota. PAR2 agonism further suppressed microbial nucleotide and vitamin B6 biosynthesis while increasing inositol degradation, TCA cycle activity, and the methylaspartate cycle, collectively pointing to altered carbon flux and cofactor metabolism in the lung microbial microenvironment. In contrast, BALF showed no significant pathway changes, consistent with its low biomass and the transient, dilution-sensitive nature of airway lumen sampling.

Sex emerged as a moderate modifier of the microbiome, with the strongest and most consistent microbial differences in the gut, followed by the lung tissue, and fewer but distinct effects in BALF. Across tissues, sex-associated taxa were dominated by *Muribaculaceae* (*Bacteroidota*) and Clostridia-associated *Bacillota A*, indicating that these lineages are particularly sensitive to sex-dependent host factors^23^.

This study has several limitations that should be considered when interpreting the findings. First, although we included 6–8 mice per condition and 3–4 mice per sex-stratified subgroup, these numbers remain modest. Our design was constrained by the requirement to use littermates and by breeding limitations; however, we minimized environmental variation by maintaining a controlled housing environment in which each cage contained a mouse from each experimental condition (WT, PAR2^-/-^, WT + HDM, PAR2^-/-^ + HDM, WT + PAR2), with three WT and two PAR2^-/-^ animals per cage. Further, because females and males were housed in separate cages, sex and cage effects cannot be fully disentangled, and some sex-associated differences may partially reflect cage-level structure despite the use of littermates. Second, we used a single intranasal challenge and an 18-hour endpoint for tissue collection. This acute timepoint was selected to capture early microbial responses to protease-rich stimuli, but it does not address temporal dynamics, compensatory microbial shifts, or the cumulative effects of chronic HDM exposure or repeated PAR2 activation that more closely model human allergic airway disease. Future studies will need to incorporate later timepoints and chronic exposure to determine whether the PAR2-dependent microbial changes observed here persist, intensify, or resolve over time. Third, both BALF and lung tissue represent low-biomass environments, which increases susceptibility to stochastic variation and may limit the detection of subtle microbial differences. Finally, we did not measure epithelial or immune responses in parallel with microbial profiling, which restricts mechanistic interpretation of how PAR2-dependent microbial changes may influence mucosal immunity.

Taken together, these findings suggest that PAR2 functions as a selective regulator of mucosal microbial ecology rather than a broad determinant of community diversity, shaping composition and predicted microbiome metabolic capacity in a compartment-specific, context-dependent, and partially sex-dependent manner. The stronger and more reproducible effects observed in gut and lung tissue, compared with BALF, also highlight the importance of sampling strategy in low-biomass airway microbiome studies and support the use of tissue-based profiling for capturing biologically meaningful host-microbe interactions. More broadly, our data position PAR2 as a potential molecular link between protease-rich environmental exposures, epithelial sensing, and microbial remodeling across mucosal surfaces. Several of the microbiome-associated patterns observed in PAR2-deficient mice resemble features reported in airway and gut disease models, including reduced microbial nucleotide biosynthesis (seen in inflamed lung tissue^24^), altered *Muribaculaceae* abundance (linked to gut barrier dysfunction^25^), and shifts in *Bacillota/Bacteroidota* balance associated with asthma susceptibility^2^. These parallels suggest that PAR2-dependent microbial states may influence disease vulnerability. Future studies should define the mechanisms underlying these PAR2-dependent shifts and determine whether PAR2-driven microbial changes actively contribute to mucosal inflammation, barrier regulation, and host susceptibility across the gut–lung axis.

## Supporting information

Fig. S1 Experimental Design

## Supplementary Table legends

**Table S1. Metadata and absolute abundance values for gut, lung, and BALF at phylum and genus levels.** Metadata for all samples (sex, genotype, treatment, tissue) and absolute abundance values for all detected phyla and genera across gut, lung, and BALF. Absolute abundances were used for all statistical analyses, including Wilcoxon rank-sum tests for genus-level comparisons and diversity analyses.

**Table S2. Relative abundance values for gut, lung, and BALF at phylum and genus levels.** Relative abundance values for all detected phyla and genera across gut, lung, and BALF. Values are expressed as the proportion of total reads per sample and used for descriptive summaries and visualization.

**Table S3. Phylum-level composition in WT controls for gut, lung, and BALF.** Mean, standard deviation (SD), sample size (N), standard error (SE), and 95% confidence intervals (CI95) for phylum-level abundances in WT samples across all three compartments.

**Table S4. Alpha- and beta-diversity metrics and statistical comparisons across gut, lung, and BALF.** Alpha-diversity metrics (Shannon, Chao1, Pielou’s evenness) with group comparisons performed using Wilcoxon rank-sum tests. Beta-diversity analyses based on Bray– Curtis dissimilarity, including overall PERMANOVA, pairwise PERMANOVA, and PERMDISP to assess dispersion. Results are reported for genotype and treatment contrasts.

**Table S5. PICRUSt2 pathway differences (sexes combined).** Significant PICRUSt2-inferred functional pathways (FDR-corrected p < 0.05) for gut and lung compartments. For each pathway, the table reports the group mean relative frequency (%), standard deviation (%), uncorrected p-value, FDR-corrected p-value, difference between group means, and the 95% lower and upper confidence interval bounds. Comparisons include WT vs PAR2^-/-^, WT + HDM vs PAR2^-/-^ + HDM, WT vs WT + HDM, and WT vs WT + PAR2 agonist. No significant pathways were detected in BALF.

**Table S6. Alpha-diversity comparisons for sex differences across gut, lung, and BALF.** Alpha-diversity metrics (Shannon, Chao1, Pielou’s evenness) compared between females and males within each genotype and treatment group using Wilcoxon rank-sum tests. Reported values include the W statistic and corresponding p-value for each comparison.

**Table S7. Significant sex-dependent differences.** Genus-level taxa showing significant female–male differences within each tissue compartment. Reported values include the comparison, sample sizes (n1, n2), Wilcoxon rank-sum statistic (W), and corresponding uncorrected p-value. Analyses were performed separately for gut, BALF, and lung using non-parametric Wilcoxon tests without multiple-testing correction.

**Table S8. PICRUSt2 pathway differences (sexes separated).** Significant PICRUSt2-inferred functional pathways (FDR-corrected p < 0.05) for gut and lung compartments analyzed separately for females and males. For each pathway, the table reports the group mean relative frequency (%), standard deviation (%), uncorrected p-value, FDR-corrected p-value, difference between group means, and the 0.5% lower and upper confidence interval bounds for the same four comparisons: WT vs PAR2^-/-^, WT + HDM vs PAR2^-/-^ + HDM, WT vs WT + HDM, and WT vs WT + PAR2 agonist. No significant pathways were detected in BALF.

## Author Contributions

Conceptualization, P.F., R.M., H.V.; methodology, M.A., R.J., R.M.; software, R.J., R.M., S.M.; formal analysis, R.J., R.M.; investigation, C.B., G.Z., M.A., M.T., R.J., R.M., S.M.; resources, P.F., M.A., H.V.; data curation, M.A., R.J.; writing—original draft, R.M.; writing—review and editing, P.F., R.J., H.V.; visualization, R.J., R.M.; supervision, P.F., H.V.; project administration, M.A.; funding acquisition, P.F., H.V. All authors have read and agreed to the published version of the manuscript.

## Acknowledgments

P.F. is the AstraZeneca (Canada) Chair in Asthma and Obstructive Lung Disease. Copilot (GenAI) was employed to generate the R code used to visualize data (**Fig. 1, 2, 3**). All code was manually inspected, edited, and executed by the authors using ggplot2 in R Studio (2024.09.0, build 375).

## Notes

### Competing Interest Statement

The authors have declared no competing interest.

### Funding

- CIHR: 0756000040

