## Supplementary figures and images for "PAR2 signaling shapes microbial and metabolic remodeling along the gut–lung axis"

### Fig. S1 Experimental Design

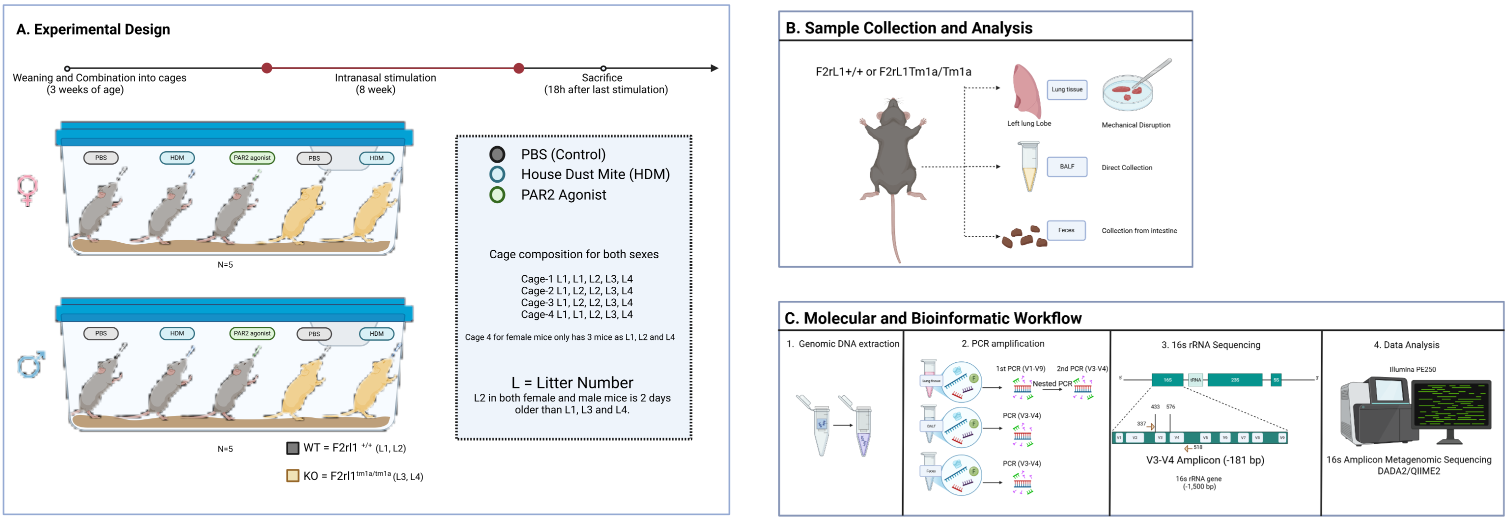
